# Interaural Time Difference Tuning in the Rat Inferior Colliculus is Predictive of Behavioral Sensitivity

**DOI:** 10.1101/2021.02.21.432131

**Authors:** Kongyan Li, Vani G. Rajendran, Ambika Prasad Mishra, Chloe H.K. Chan, Jan W. H. Schnupp

## Abstract

Recent studies have shown that rats are a useful model for binaural cochlear implant (CI) research, with behavioral sensitivity to interaural time differences (ITDs) of CI stimuli which are better than those of human patients. Here, we characterize ITD tuning in the rat inferior colliculus (IC) and explore whether quality of tuning can predict behavioral performance. We recorded IC responses to stimuli of varying pulse rates and envelope types and quantified both mutual information (MI) and neural *d*′ as measures of ITD sensitivity. Neural *d*′ values paralleled behavioral ones, declining with increasing click rates or when envelopes changed from rectangular to Hanning windows. While MI values increased with experience, neural *d*′ did not. However, neural *d*′ values correlated much better with behavioral performance than MI. Thus, neural *d*′ appears to be a particularly well suited to predicting how stimulus parameters will impact behavioral performance.

## Introduction

Cochlear implant (CI) users face enormous challenges when trying to localize sound. The inability of CI users to use interaural time difference (ITD) cues in particular has been thought be due to the lack of normal hearing experience during a critical period in development [1-3]. However, new evidence is beginning to challenge this theory. Specifically, neonatally deafened, adult rats fitted with bilateral CIs that are precisely synchronized to provide ITD cues in the normal physiological range were able to learn to lateralize CI pulses with ITD thresholds that were indistinguishable from those normal hearing animals [4], in contrast to prelingually deaf human CI users, who often have poor ITD thresholds [5]. These findings raise the possibility that the reasons for poor ITD discrimination in current bilateral CI users may be more to do with technological shortcomings in devices in current clinical use which were not from the outset designed with optimal binaural stimulation in mind.

Two parameters of particular relevance to CI processing strategies are pulse rate and envelope shape, both of which influence the detectability of ITDs of pulse train stimuli. A recent study explored these parameters with acoustic pulse train stimuli in normal hearing rats and found robust behavioral sensitivity to envelope ITD that declined for high pulse rates (≥ 900 Hz) and were poorer for Hanning windowed stimuli than for rectangular windowed stimuli [6]. In this context, studying the neural tuning to ITDs of pulse train stimuli in the rat brain merits further exploration, given that all CIs in current clinical use encode sounds as electrical stimulus pulse trains. Since acute electrophysiological experiments on anesthetized animals are often quicker and easier to perform than animal psychoacoustics studies, an important question for translational research is whether measures of the quality of ITD tuning recorded along the auditory pathway can serve as good predictors of psychoacoustic behavioral performance limits. To take a first step towards answering this question, we focus here on 2-alternative forced choice (2afc) lateralization performance, while acknowledging that free field localization is a different, harder problem which is beyond the scope of this study. In the context of 2afc lateralization, *d*′ is an established way to compute behavioral performance limits. If we capture the theoretical “lateralization ability” of a neuron through *d*′, an obvious question becomes whether changes in neural *d*′ with varying stimulus conditions mirror corresponding changes in behavioral *d*′. If so, measuring how various novel binaural CI stimulation strategies affect neural *d*′ could provide a relatively simple test to decide which strategies may be most promising for further research and development.

Mutual information (MI) is another very widespread method for quantifying the quality of neural coding. Unlike a *d*′ analysis, MI does not explicitly quantify how well neural responses lateralize sound sources, but merely how much of the variability (“entropy”) of the responses can be accounted for by variation in the stimuli. Since MI was not tailor made for discrimination tasks we might intuitively expect changes in MI under various stimulus conditions to correlate less well with psychoacoustic lateralization performance than neural *d*′, but to our knowledge this has not been explicitly tested.

To explore these questions, we recorded extracellular analog multiunit activity in the central nucleus of the inferior colliculus (IC) of anesthetized, normal hearing rats. The IC is a suitable target for recordings as it is a much studies obligatory relay nucleus of the ascending auditory pathway, and recent work highlights its importance in ecologically important auditory processing tasks such as binaural unmasking [7, 8]. ITD tuning curves were recorded with pulse train stimuli with rates of 50, 300, 900, 1800 or 4800 Hz, with Hanning or rectangular envelopes, were tested on two cohorts of animals, one naive and one that had been previously trained on a 2afc sound lateralization task using the same stimuli. Our study had three aims: 1) to compute a neural *d*′ metric that estimates the theoretical ability for a neuron to discriminate left-leading from right-leading ITDs, 2) to explore the dependence of neural *d*′ and MI on pulse rate, envelope shape, and prior training on a sound lateralization task, and 3) to compare the usefulness of neural *d*′ or MI as predictors of behavioral sensitivity to ITD.

## Materials and Methods

### Animals

Eleven female Wistar rats were used in this study: Six rats aged 8 weeks without any prior behavioral training formed the “naive” cohort, while five mature rats aged 76 to 86 weeks who had received extensive training and testing in sound lateralization [6] formed the “experienced” cohort. We label the cohorts as “naive” vs “experienced” to acknowledge that the cohorts differ substantially in age as well as in training, with our naive rats being young adults, and the experienced animals being what might be called “late middle age”. In the results we will see that the neural tuning in the two cohorts was very similar, but where there were significant differences these are most easily interpreted in terms of experience dependent increases stimulus related information coding rather than specific training effects or age related declines. Rats were housed in 12/12 h light/dark cycle with 2 or 3 animals in one cage and *ad-lib* food.

### Extra-cellular recording

#### Stimuli

In the physiological experiments described here, 100 ms click trains enveloped with a rectangular or Hanning window were delivered to the ear canals of the anesthetized animals as described below. Pulse rates were 50, 300, 900, 1800 or 4800 Hz, and stimuli were delivered at an average binaural level of 80-85 dB SPL, at a sampling rate of 48828.125 Hz. The stimulus to one ear was offset from that in the other by an integer numbers of samples to generate ITDs varying between -164 μs (left ear leading) and +164 μs (right ear leading) in 20.5 μs steps.

Stimuli were generated by an RZ6 multi-I/O processor (Tucker-Davis Technologies, USA) and delivered over a pair of custom-made earphones based on Knowles ED-27288-000 miniature speakers connected to stainless steel hollow stereotactic ear bars. These were placed inside the ear canals of the rat and served to fix the rat into a stereotaxic frame (RWD Life Sciences, China). The earphone assemblies were calibrated using a G.R.A.S. 46DP-1 microphone.

#### Surgical Procedure

Rats were anesthetized by intraperitoneal injection of ketamine (80mg/kg, 10%, Alfasan International B.V., Holland) and xylazine (12 mg/kg, 2%, Alfasan International B.V., Holland). The integrity of the rats’ tympanic membranes and outer ear canals were checked visually, and acoustic brainstem responses (ABRs) were recorded to confirm that the animals had normal hearing thresholds. The rats’ eyes were rinsed with 0.9% Sodium Chloride (Normal Saline, B.Braun Medical Industries Sdn. Bhd, Malaysia) and eye gel was applied (Lubrithal, Dechra Veterinary Products A/S Mekuvej 9 DK-7171 Uldum) to prevent drying.

Following ABR recording, the analgesic Carprofen (5 mg/kg, 50 mg/ml, Norbrook Laboratories Australia Pty Limited, UK) was subcutaneously injected. The scalp was shaved and an incision was made along the mid-line to expose the skull. Lignocaine (0.3 ml, 20 mg/ml, Troy Laboratories Pty.Limited, Australia) was applied to the incision and the surface of the skull for additional local anaesthesia. A craniotomy was then made to expose the right occipito-parietal cortex, from the sagittal suture, extending roughly 3 mm rostral and 1 mm caudal to lambda and 4 mm lateral from mid-line. The absence of a toe pinch reflex was checked periodically to monitor anesthesia depth, and, if necessary, anesthetic was supplemented by i.p. injection of one third of the initial anaesthesia induction dose. Atropine sulphate (0.13 mg/kg, 0.65 mg/ml, Troy Laboratories Pty. Limited, Australia) was given i.p. if the rat’s heart rate was slow. During the experiment, anesthesia was maintained by continuous i.p. infusion of a 0.9% sodium chloride solution of ketamine (17.8 mg/kg/h, 10%, Alfasan International B.V., Holland) and xylazine (2.7 mg/kg/h, 2%, Alfasan International B.V., Holland) through a syringe pump running at a rate of 3.1 ml/h.

#### Electrophysiological Recording

Neural signals were captured through a PZ5 neurodigitizer (Tucker-Davis Technologies, USA) at a 24414.0625 Hz sample rate and processed via RZ2 bioamp processor (Tucker-Davis Technologies, USA). Extracellular multiunit activity was record using a 32-channel single shaft electrode (E32-50-S1-L6, ATLAS Neuroengineering, Belgium) inserted vertically into the rat’s right inferior colliculus (IC) to a depth of approximately 5.0 mm below the surface of the occipital cortex, using a micromanipulator (RWD Life Sciences, China). A white noise burst search stimulus was presented, and robust, short latency (3-5 ms) responses to the sounds were taken as an indication that recording sites were likely inside the central nucleus of the IC. At the beginning of every penetration, single click stimuli at various sound levels were presented to the left (contralateral to the recording side) ear to estimate neural response thresholds, and then the test stimuli were presented. At the end of the experiment, animals were overdosed with pentobarbital sodium (1∼2 ml, 20%, Alfasan International B.V., Holland).

### Analysis of electrophysiological data

#### Data preprocessing

Analog multiunit activity (AMUA) was calculated following the procedure used in [9]. First, traces were bandpass filtered between 300 Hz and 6000 Hz using a 3^rd^ order Butterworth filter. The absolute value was then taken, and then a low pass 3^rd^ order Butterworth filter was applied below 6000 Hz. In this study, traces were additionally downsampled by a factor of 20 using the Matlab function ‘decimate()’. The resulting AMUA traces used for subsequent analysis were thus at a sampling rate of 1220.7 Hz.

Stimuli were 100 ms in duration and were followed by at least 400 ms of silence before the next stimulus was delivered. Baseline-corrected neural responses to each stimulus were calculated by subtracting baseline activity (average AMUA in a 150 ms window, 155-305 ms post stimulus onset) from the stimulus-driven “onset response” (average AMUA in a 50 ms window, 5-55 ms post stimulus onset) in each trial. Baseline-corrected neural responses were used in all subsequent analyses. To verify that the results presented here are not very sensitive to the choice of the response window, all analyses described below were also repeated with a “sustained” response window from 5-105 ms post stimulus onset. This yielded very similar results to those presented below for the 50 ms response window, even if mutual information (MI) values were smaller overall for the longer response windows.

#### Mutual information calculation

To determine which recorded units were significantly tuned to ITD, and to quantify the strength of tuning, we computed the MI between the 17 tested ITD values and the neural response. Since we might expect MI values to be low for stimulus conditions that had been found to be behaviorally challenging (e.g. 4800 Hz pulse rate), a relatively generous inclusion criterion was set: a unit was included in subsequent analyses if it showed significant MI for any of the ten (5 pulse rates x 2 envelopes) stimulus conditions tested. MI values were calculated using the adaptive direct method described in [10], using 5 initial discretization levels for the neural response values. Raw MI values were bias corrected by subtracting the average MI value obtained after scrambling which ITD condition each trial belonged to 100 times. Following the convention established in [10], units were considered to have significant MI for ITD if they satisfied two conditions: their raw MI was larger than their bias estimate for 100 of the 100 iterations (p < 0.01), and the raw MI was at least twice as large as the bias estimate. Only multiunits that satisfied both of these conditions (925 out of 1280 multiunits) were considered significantly ITD sensitive.

#### Calculation of neural *d*′ based on ROC analysis of ITD tuning curves

The rationale behind the ROC analysis was to estimate each recorded multiunit’s “ability to discriminate left from right” in the sense of whether the delivered stimuli had a “positive” (right-ear-leading) or “negative” (left-ear-leading) ITD. The resulting metric is a *d*′ value that can be compared to *d*′ values obtained behaviorally. The method used here is inspired by a similar approach described in [11] to quantify the neural sensitivity to a stimulus parameter.

Neurons in the IC of small mammals are well known to show predominantly contralateral spatial tuning, meaning that stronger responses are typically expected for ITDs leading in the ear opposite to the side of the brain where recordings are made. For ROC analysis, first, for each given envelope and click rate, trial-by-trial neural responses for all contralateral ITD stimulus presentations (8 contralateral ITD values × 30 repeats = 240 values) and all ipsilateral ITD stimulus presentations (8 ipsilateral ITD values × 30 repeats = 240 values) were assembled. An ROC curve was constructed by considering a range of decision criteria above which the neural response would indicate that the stimulus had a contralateral ITD. The “hit rate” was defined as the proportion of trials with neural responses above the decision criterion that actually came from a contralateral ITD stimulus, and the “false alarm” rate was the proportion of trials with neural responses above the decision criterion that came from an ipsilateral ITD stimulus. The ROC curve plots the hit rate as a function of the false alarm rate as the decision criterion is moved across the full range of the response distribution from both positive and negative ITD trials. The area under the ROC curve (AUC) ranges between 0 and 1, and corresponds to the probability of correct responses in a 2afc task. It is therefore directly related to *d*′ (Equation 1), which is calculated by multiplying the square root of 2 by the inverse of the standard normal cumulative distribution function *Z* (the function ‘norminv()’ in Matlab) evaluated at the probability value (Z(*p*), *p* ∈ [0,1]) specified by the AUC.

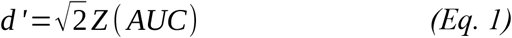

The resulting *d*′ values are interpretable as a multi-unit’s ability to inform lateralization judgments on the incoming stimuli. AUC values close to 0.5 result in *d*′ values near zero and are indicative of performance at chance. Contralaterally tuned units (i.e. those preferring stimuli with contralateral-ear-leading ITDs) will yield AUC values > 0.5 and positive *d*′ values, while units that are tuned ipsilaterally will yield negative *d*′ values. Larger absolute *d*′ values (|*d*′|) thus indicate higher ITD sensitivity irrespective of whether a unit happens to be contralaterally or ipsilaterally tuned.

#### Behavioral training (for the experienced cohort)

The 5 experienced animals had previously been subjects of our previously published behavior study [6], and their psychoacoustic training and testing has been described in detail there. Briefly, the rats were trained 5 days per week, using drinking water as positive reinforcer in an instrumental conditioning task. Rats were trained to perform 2afc lateralization of acoustic pulse trains which had either a left-leading or right-leading ITDs. In a custom behavioral cage with three spouts, the animals initiated trials by licking the center spout, and indicated whether they heard click train stimuli as coming from the left or right by licking the corresponding response spouts. Correct responses (on the side of the leading ear) were positively reinforced with a few drops of drinking water, incorrect responses triggered a “timeout” period of 15-30 s during which no new trials could be initiated and a negative feedback sound was played. Rats performed 2 training sessions per day, with each session last about 30 min, resulting in ∼200 trials per day. Stimuli were 200 ms broadband click trains with either a rectangular or a Hanning window envelope and pulse rates of 50, 300, 900, 1800, 2400 or 4800 Hz. The sounds were presented at a sample rate of 44,100 Hz and an average binaural acoustic level of approximately 80-85 dB SPL. The ITD values tested spanned the physiological range of the rat, ranging from 158.9 μs left-ear-leading to 158.9 μs right-ear-leading in 22.7 μs steps. ITDs varied randomly from trial to trial, and in each session a subset of the aforementioned click rates and window types were tested.

#### Behavioral *d*′ analysis

To be able to compute behavioral *d*′ values to compare against the neural *d*′ analysis described above, we reanalyzed the data collected during the [6] study as follows. For each animal, we pooled data across all behavior testing sessions, and defined as “hit rate” (HT) the proportion of trials with stimulus ITD > 0 during which the animal responded on the right, averaged for all right-ear-leading ITD values. Similarly, we defined the “false alarm rate” (FA) as the proportion of trials with left-ear-leading ITD for which the animal responded on the right. We then simply inserted these values into the well-known formula for behavioral *d*′ values for 2afc tasks; *d’ = z(HT) – z(FA)*, where *z* again stands for the inverse cumulative Gaussian normal distribution. Because the range of ITDs tested in each case was very similar (+/-164 μs in 20.5 μs steps for the electrophysiological data, +/-159 μs in 22.7 μs steps for the behavior data), the overall “task difficulty” was also comparable in the two conditions, making it reasonable to compare neural and behavioral *d*′ values directly.

#### Statistical analysis: mixed effects ANOVA

It is well know that IC neurons located in close anatomical proximity tend to have similar tuning to acoustic stimuli, and consequently multiunit responses recorded from adjacent recording sites along a single multielectrode penetration are likely to be correlated, and cannot be treated as statistically independent samples. To determine whether the stimulus conditions tested (click rate, envelope, experience) had main effects or interaction effects on |*d*′|, while compensating for the lack of statistical independence between channels recorded from the same penetration, a mixed effects analysis of variance analysis (ANOVA) was used, which used penetration number as a random effect. Given that ANOVA assumes normally distributed data, the test was run on values of |*d*′| which were monotonically transformed by taking the cube root in order for the transformed values to be approximately normally distributed,. The mixed effects ANOVA model comprised the following terms: an intercept (1), all main effects (3), 2-way (3), and 3-way (1) interaction effects for a total of 8 parameters or explanatory variables.

## Results

### Roughly 72% of recorded IC multiunits showed significant mutual information for ITD

A first analysis on all recorded multiunit channels was performed to check for ITD tuning using the well-established metric of mutual information. Since our stimuli tested a wide range of pulse rates with Hanning and rectangular windows, we expected that some conditions, for example the 4800 Hz pulse rate, which animals were unable to lateralize behaviorally, were also less likely to produce ITD tuned responses in the IC. Thus, for a multiunit to be included in further analysis, it needed to have a bias-corrected MI value significantly greater than zero (p<0.01, permutation test) and this bias-corrected MI needed to be at least twice as large as the bias to avoid false positives (see *Methods*). Since MI was computed for each unit across ten possible stimulus conditions (5 pulse rates x 2 envelopes), we selected for analysis all multiunits that could satisfy the above two criteria in at least one of the ten conditions tested. Based on this, we found that 925 of the 1280 recorded multiunits showed significant sensitivity to ITD based on mutual information (Fig. 1).

**Fig 1.**
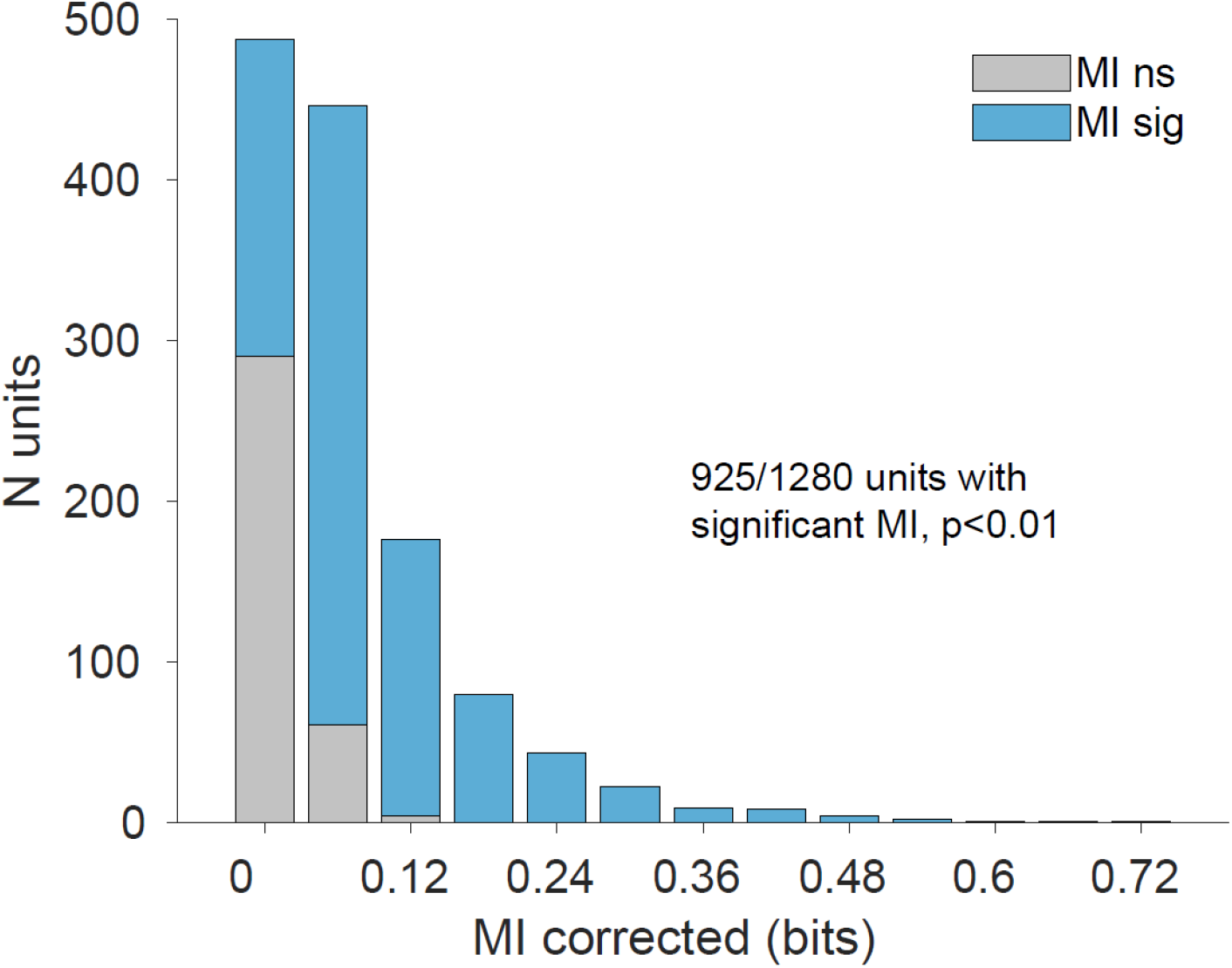
Distribution of mutual information (MI). Of the 1280 recorded multiunits, 925 (∼72%) showed bias-corrected MI greater than zero (p<0.05) and a corrected MI value greater than twice the bias for at least one of the pulse rate x envelope conditions tested. Subsequent analyses are based on these 925 multiunits.

### Neural |*d*′| and MI values are often, but not always, correlated

To explore the ability of these neurons to lateralize ITDs based on response strength, a neural *d*′ sensitivity index was computed using a receiver operating characteristic (ROC) analysis (see *Methods*). Briefly, a multiunit’s trial-by-trial responses for all contralateral ITD trials of a given condition were pooled together, and the same was done for all ipsilateral ITD trials. A small overlap between contra ITD and ipsi ITD response distributions (Fig. 2B) would indicate that a multiunit is highly sensitive to whether sound comes from the left or right. This is captured by the ROC curves shown in Fig. 2C, which plot hit rates against false alarm rates for a decision threshold that is moved across the range of the data in Fig. 2B. A *d*′ value can then be calculated directly from the area under the ROC curve (see *Methods*). Multiunit responses often resulted in MI and *d*′ values that were either both high or both low (first and fourth rows of Fig. 2, respectively). However, this need not necessarily be the case, for example if tuning curves are somewhat symmetric about 0 ITD, which could result in a large MI but a low *d*′ (Fig. 2, second row). The third row of Fig. 2C shows a contrasting example with high *d*′ and a low MI. Note that a *d*′ of 1 indicates that an ideal observer should be able to use the multiunit’s responses to lateralize ITDs with over 75% accuracy. (This follows from eqn 1, noting that the cumulative normal distribution value of 1/√2 ∼ 0.76). Similarly, a *d*′ of 2.24 (Fig. 2C, top panel) corresponds to nearly 95% accuracy, and a *d*′ of 0.81 (Fig. 2C, third row) corresponds to roughly 71% accuracy.

**Fig 2.**
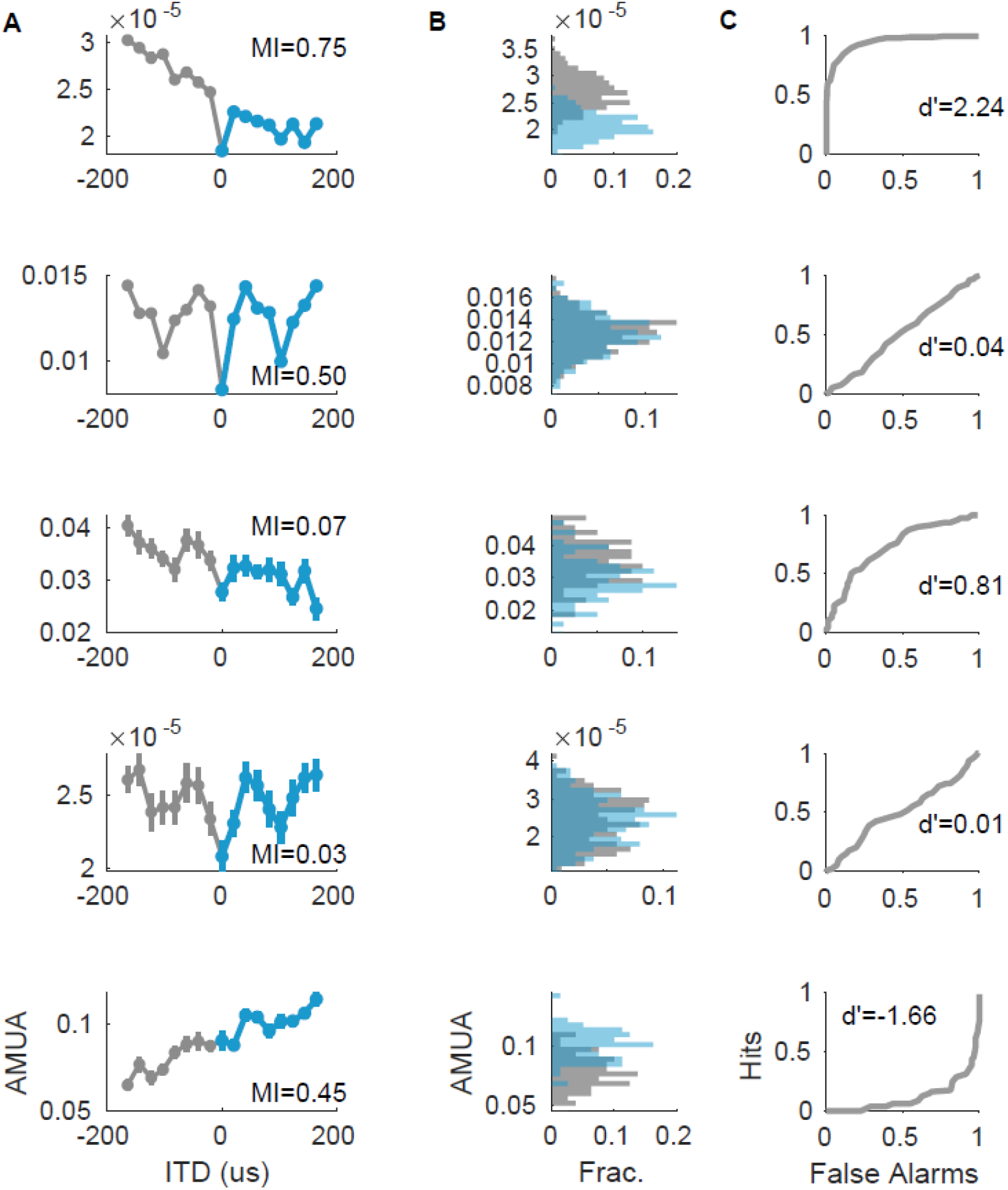
Calculation of neural d-prime and a comparison with MI for example multiunits. 1^st^ row – high MI, high *d*′; 2^nd^ row – high MI, low *d*′; 3^rd^ row – low MI, high *d*′; 4^th^ row – low MI, low *d*′; 5^th^ row – negative *d*′ due to ipsilateral tuning. **A**) Tuning curves showing mean +/-SEM of neural responses as a function of stimulus ITD. **B**) Histograms pooling trial-by-trial neural responses for all ipsilateral (blue) or contralateral (gray) ITD conditions. **C**) ROC curve based on splitting the AMUA range in panel B into 50 linearly spaced ‘criterion’ points. Hits were defined as the proportion of contralateral ITD responses larger than a given criterion value, and false alarms were defined as the proportion of ipsilateral ITD responses larger than the given criterion value. Note the correspondence between the degree of overlap in panel B and the resulting *d*′ value.

Generally, while *d*′ values capture an upper bound on the ability to use the strength of a multiunit’s response to distinguish contralateral from ipsilateral ITDs, the MI quantifies an “entropy reduction” in the uncertainty about which of any of the 17 possible ITD values was presented at a given trial, and it tends to be high when trial-to-trial variability is low compared to the difference in the average response between any pair of ITD values, regardless of whether these differences are systematically distributed across the midline or not.

Only a small minority of multiunits with significant MI (N=20, ∼3% of total) had negative *d*′ values due to ipsilateral tuning (Fig. 2, fifth row). The analyses that follow report on |*d*′| since the magnitude of *d*′ reflects discriminability of neural responses to left and right ITDs regardless of contralateral or ipsilateral tuning.

### Both MI and |*d*′| depend on pulse rate and envelope, but MI is affected by experience while |*d*′| is not

With neural |*d*′| as an additional quantification of ITD tuning, we now turn to the question of how ITD tuning varies with click rate and envelope, two important parameters in cochlear implant stimulation. Additionally, by comparing |*d*′| values from the naive and the experienced cohort, we could also examine the effect of experience on ITD tuning in the IC. A mixed effects ANOVA was performed, treating penetration ID as a random effect since multiunits from the same penetration cannot be considered independent of each other. As shown in Table 1, mutual information shows main effects of all three experimental parameters. Click rate, envelope, and experience all affect the mutual information between IC responses and ITD. Likewise, neural |*d*′| is also affected by click rate and envelope, but interestingly, not by experience (Table 2).

**Table 1:** ANOVA of effects of stimulus parameters and experience on MI. Fixed effects coefficients (with 95% confidence intervals) returned by a mixed effects ANOVA. MI is significantly affected by pulse rate, envelope and experience.

| Name | Estimate | Lower | Upper | SE | tStat | DF | pValue |
| --- | --- | --- | --- | --- | --- | --- | --- |
| (Intercept) | 0.3816 | 0.3136 | 0.4495 | 0.0347 | 11.0100 | 9246 | 0.00000 |
| Pulse Rate | -0.0041 | -0.0059 | -0.0023 | 0.0009 | -4.4802 | 9246 | **0.00001** |
| Envelope | 0.0105 | 0.0054 | 0.0156 | 0.0026 | 4.0626 | 9246 | **0.00005** |
| Experience | 0.0516 | 0.0145 | 0.0886 | 0.0189 | 2.7287 | 9246 | **0.00637** |

**Table 2:**
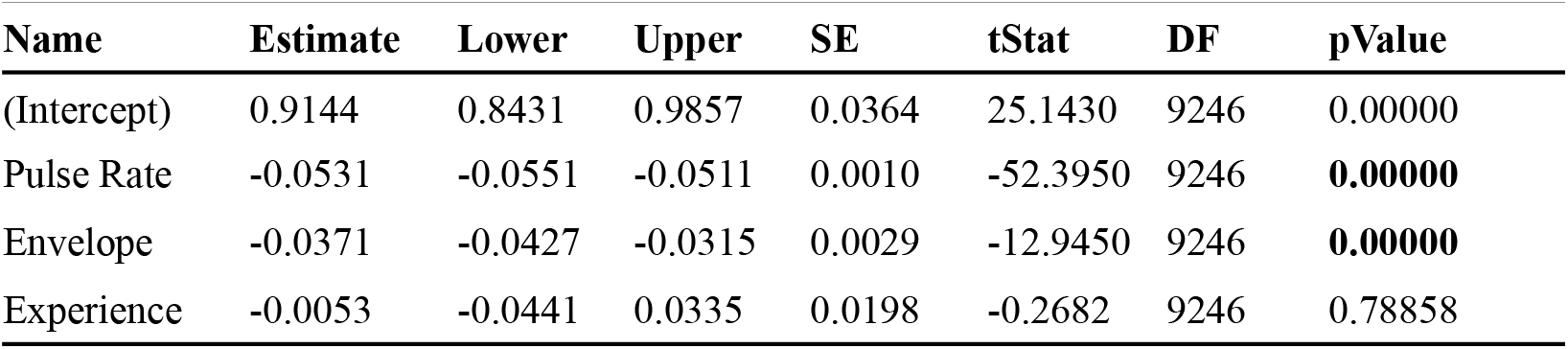
ANOVA of stimulus parameters and experience on neural |*d*′ |. Fixed effects coefficients (with 95% confidence intervals) returned by a mixed effects ANOVA. Neural |*d*′| shows main effects of pulse rate and envelope, but not experience.

Fig. 3 shows the distribution of MI and |*d*′| values for each of the conditions tested. For neural |*d*′| (Fig. 3A-B), values tend to decline with increasing pulse rate, dropping dramatically at and above 900 Hz, for Hanning window stimuli in particular. There is no obvious difference in the distributions of neural |*d*′| between experienced and naive animals, but MI (Fig. 3C-D) is higher in experienced animals. Population |*d*′| tended to be highest at 300 Hz and decline to near zero at high click rates. Neural |*d*′| values were also consistently larger for rectangular window stimuli than for Hanning window stimuli. Notably, at low pulse rates, a number of multiunits show |*d*′| values well above 1, indicating that these multiunits could theoretically distinguish left and right ITDs with well over 75% accuracy. At 300 Hz, the best multiunits had |*d*′| at or above 2, and the median |*d*′| of the entire neural population is around 0.5, corresponding to a median accuracy in discriminating left and right of ∼64% for individual multiunits. At 900 and 1800 Hz, there is a substantial drop in the IC’s ability to distinguish left ITDs from right, with a slight advantage for rectangular windowed stimuli. At 4800 Hz, the IC appears equally poor in distinguishing left from right for both Hanning and rectangular windowed stimuli, and even the largest |*d*′| values don’t reach beyond around 0.5.

**Fig 3.**
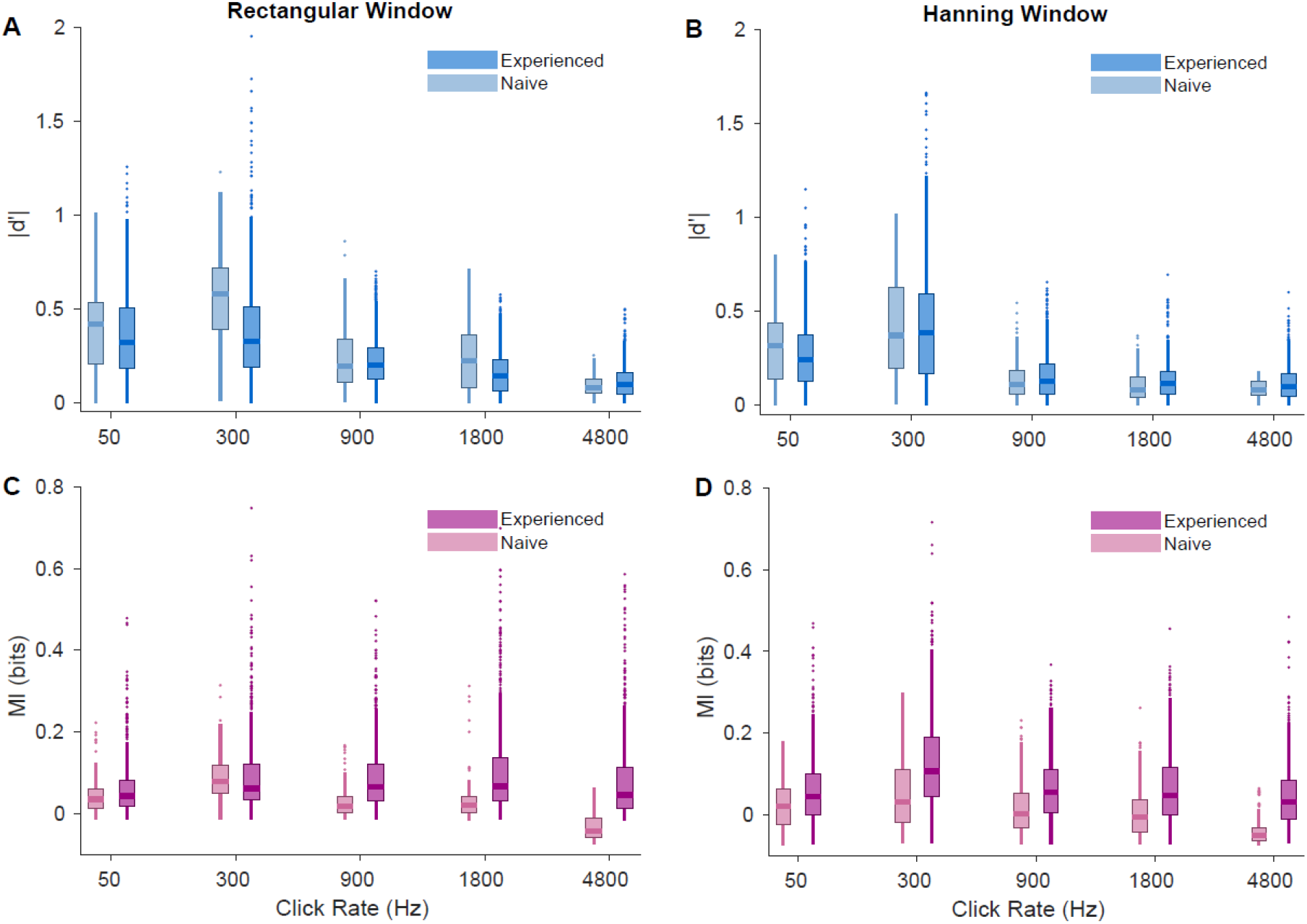
Neural |*d*′| and MI values for all conditions tested. **A)** Neural |*d*′| for rectangular windowed stimuli. **B)** Neural |*d*′| for Hanning windowed stimuli. **C)** MI for rectangular windowed stimuli. **D)** MI for Hanning windowed stimuli.

### Neural |*d*′| is highly correlated with behavioral *d*′ measured in a 2afc sound lateralization task

Having established that an IC neuron’s ability to encode ITD is affected strongly by pulse rate and envelope shape, we now turn to the question whether these variations in the quality of ITD tuning in the IC are predictive of similar variations in behavioral ITD lateralization performance. Fig. 4A-B summarizes behavioral data for the five experienced animals alongside the neural |*d*′| and MI values recorded from the same animals. To summarize the distributions shown in Fig. 3 with a single representative value in Fig. 4 we chose to plot the 95th percentiles of the distributions for MI (MI_95_) and neural |*d*′| (|*d*′|_95_) at each stimulus condition and for each animal, reasoning that the 95^th^ centile would be representative of multiunits with “strong tuning” without being very sensitive to outliers. Behavioral *d*′ is often substantially larger than neural |*d*′|_*95*_, but both show parallel declines for increasing stimulus pulse rate. Such parallel trends are not apparent when comparing behavioral *d*′ with MI_95_.

**Fig 4.**
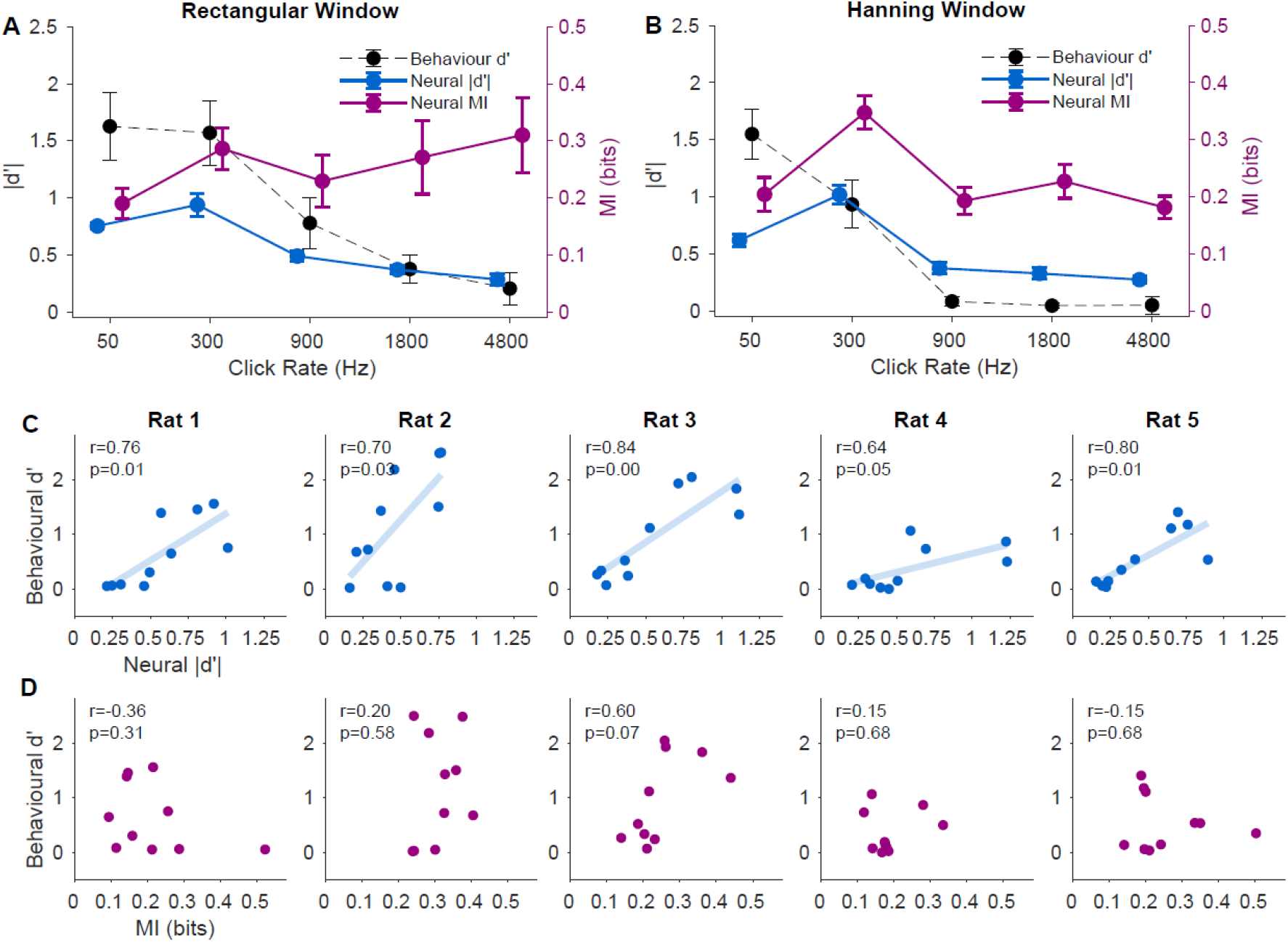
Comparison of neural and behavioral ITD sensitivity, Neural |*d*′| (blue) co-varies with behavioral *d*′ (black), but MI (violet) does not. **A, B)** the 95^th^ centile of the distribution of multiunit neural |*d*′| (|*d*′|95) and MI values (MI95) were computed for each animal at each stimulus condition, and means (+/-SEM) across animals are plotted against stimulus pulse rate. For comparison, across-animal mean (+/-SEM) behavioral *d*′ values for the same stimulus conditions are shown in black. **A)** For rectangular windowed stimuli, the gradual decline in neural |*d*′|95 with increasing stimulus click rate is reflected in a parallel downward trend in behavioral *d*′ values. MI95 in contrast shows no such systematic decline. Instead, variability in MI95 between animals increases with pulse rate. **B)** For Hanning windowed stimuli, there is a marked drop in neural |*d*′| for pulse rates ≥900 Hz that is also reflected by parallel declines in behavioral *d*′. This trend is less apparent for MI95. **C)** Neural |*d*′|95 is strongly correlated with behavioral *d*′ for all five experienced animals. **D)** Neural MI95 is not significantly correlated with behavioral *d*′ for any of the five experienced animals.

Fig. 4C-D illustrate the relationship between behavioral *d*′ in each condition and neural ITD tuning for each animal in the experienced cohort. For all five animals tested, behavioral *d*′ correlated strongly with neural |*d*′|_95_. In contrast, MI_95_ did not significantly correlate with behavioral performance for any of the animals. Interestingly, individual differences in recorded neural |*d*′|_95_ values were relatively smaller than individual differences in psychoacoustic performance, with Rat 2 showing particularly high maximal behavioral *d*′, followed by Rats 1 and 3, then finally Rat 4 and 5. Note also that the behavioral *d*′ values are usually larger than the neural |*d*′|_95_ values, but this difference is more pronounced in the animals with the highest psychoacoustic performance. This leads to different regression slopes in Fig. 4C, and suggests that the neural encoding of ITDs at the level of the midbrain may not have been the main factor limiting psychoacoustic performance. Animals likely have to combine information from groups of multiunits appropriately to achieve high behavior scores, and the efficiency of these downstream read-out processes may be more of a limiting factor in determining individual psychoacoustic performance than the accuracy of the neural code at the level of the midbrain. Nonetheless, changes in neural |*d*′| as a function of changing stimulus conditions are reflected in corresponding changes in behavioral performance in all animals tested, and neural |*d*′| does not change significantly with experience. Consequently, measuring the effect of stimulus manipulations on neural |*d*′| in naive animals, for example in the context of developing new assistive or prosthetic technologies, is likely to be a useful predictor of how these manipulations will affect behavioral discrimination performance.

## Discussion

In this study, we have investigated the dependence of neural ITD coding on stimulus parameters such as pulse rate and stimulus envelope using two quite different methods for quantifying neural coding: MI and *d*′. We have compared MI and neural *d*′ against each other, as well as against behavioral *d*′ data obtained under the same stimulus conditions in some of the same animals. Our key findings can be summarized as follows:

1. MI and neural |*d*′| often, but not always, correlate, and where they differ neural |*d*′| is a better predictor of behavioral performance in a 2afc behavioral task. Curiously, MI increased with experience but |*d*′| did not.
2. Both neural and behavioral *d*′ showed similar dependence on pulse train envelope and rate, with notable declines in ITD sensitivity at high pulse rates and soft, Hanning as opposed to sharp, rectangular pulse train envelopes. In each of 5 animals for which both behavioral and neural *d*′ data were available, these two correlated strongly, indicating that changes in neural *d*′ following a given change in stimulus parameters are likely to be a useful indicator of how these changes will affect psychoacoustic performance.

These two main findings have a number of implications, both for fundamental neuroscience (which method for quantifying neural sensitivity is most appropriate under which circumstances?) and for translational research (how might stimulus processing strategies in CIs be improved to achieve better binaural hearing for CI users?) We shall briefly discuss each of these issues in turn.

### Comparing and contrasting *d*′ and MI metrics

As was already described in the context of Fig. 2, our neural *d*′ analysis was set up specifically to make a binary discrimination between any contralateral from any ipsilateral-ear-leading ITD, and it is therefore insensitive to response strength differences that occur only within one side of auditory space. In contrast, the MI analysis quantifies the discriminability between any pair of ITD values, irrespective of whether they span the midline or not. It is therefore perhaps not entirely surprising that MI and *d*′ values are not always correlated, and that neural |*d*′| values, which were computed specifically to quantify what a neural rate code might contribute to a 2afc lateralization judgments, follow trends in behavioral lateralization performance better than “general purpose” MI values which were computed without first categorizing stimuli as left-ear or right-ear-leading. Our MI measure would probably have correlated better with behavioral performance if the behavior had tested free-field localization with many more than two possible choices, but this will require confirmation in future studies. Meanwhile, the results of our study support the common sense notion that neural decoding needs to be appropriately matched to the particulars of the behavioral task one is trying to elucidate, and popular general purpose metrics including not just MI but also neural “signal-to-noise-ratios” or “signal-to-total-variance-ratios” (e.g. [2]) may be less suitable for predicting likely behavioral outcomes.

Perhaps related to this notion is the following observation: a popular account for neural mechanisms of binaural hearing is the popular “opponent channel model” [12-14] which envisages a representation of azimuthal sound source location through two populations of binaural neurons which are tuned locations in the contralateral hemifield -in other words their tuning curves would resemble the one shown in the top row of Fig. 2 are very much the rule. However, for this type of tuning curve, MI and *d*′ would be correlated, and should therefore both correlate with behavior. The observation in Fig. 4 that MI and *d*′ differ substantially in how well they correlate with performance in this 2afc lateralization task thus serves to remind us that, while hemifield tuning is the rule in ascending auditory pathway, there are also plenty of exceptions, and the representation of ITD across the population of IC neurons is more complicated than a simple opponent channel model would suggest [15].

This also leads us to a perhaps perplexing observation. On the one hand, the data in our study clearly show that neural *d*′ does not increase with experience but does correlate with perceptual performance – in other words, ITD coding in the IC of naive, young adult rats is already as good as it needs to be to support high levels of performance in behavioral ITD lateralization, and it does not increase significantly with experience. On the other hand, perhaps surprisingly, MI values which do not correlate with behavior, are nevertheless higher in the experienced animals. That neural MI values in the central auditory pathway can increase with experience is not surprising in itself, and has been described previously (e.g. [16]). What may appear counter-intuitive is that we see increases in MI even though MI is a poor predictor of behavioral performance in the 2afc task. However, several theoretical papers [17, 18] have shown that increases in MI between stimuli and neural responses can occur purely from the application of local, unsupervised synaptic learning rules in neural networks. Supervised learning with reinforcement signals may therefore not be required to increase MI, and changes in MI could in such cases occur quite independently of training in a specific task. We should therefore not be too surprised if the MI between neural firing and sound ITD increases with experience, even if the observed increases in MI appear unrelated to the reinforcement criteria chosen for a particular 2afc lateralization task which formed a part of the animals’ experience.

### The use of neural |*d*′| in translational research on binaural hearing

In this study, we described how neural encoding of ITD of pulse train stimuli in rat IC depended on pulse rate and envelope, and showed that quantifying this encoding using neural |*d*′| yields a measure that which is highly predictive of behavioral *d*′. Specifically, neural |*d*′| follows behavioral *d*′ in that it is lower for Hanning windowed stimuli than for rectangular windowed stimuli, and that it drops dramatically for pulse rates at and above 900 Hz. Note that, in this study, behavioral data were collected separately from physiological recordings, and recordings were taken from anesthetized animals. No doubt, there are well documented effects of changes in an animal’s attention in neural responses in IC [19] which are probably mediated by corticofugal backprojections [20], but while such rapid, dynamic effects can modulate midbrain tuning, they do not need to be engaged to reveal baseline neural tuning properties which do place constraints on psychoacoustic performance. If that was not the case, the clear parallels between neural and behavioral *d*′ observed in our study would be very difficult to explain. Criticisms of the use of anesthetized preparations for auditory work are often based on studies which lack control data, while studies which have directly compared responses of central auditory neurons in awake and anesthetized recordings (e.g. [21]) found little difference in neural coding. The fact that much of value can still be learned from acute, anesthetized auditory physiology experiments is reassuring because such experiments are much easier to perform quickly and at scale than experiments requiring elaborate and lengthy awake recording and behavioral conditioning protocols. As mentioned in the introduction, our work is in part motivated by a clinical need to improve binaural hearing outcomes for hearing impaired patients. We shall therefore now turn to discussing how our results relate to previous work on binaural work in humans and in CI recipients, and how our approach may contribute to future research into improved CI processing strategies.

Both the physiological data presented here and our behavioral results [6] show very similar declines in ITD stimuli for high stimulus pulse rates. Similar declines in ITD sensitivity with pulse and AM rates above 500 Hz have been previously reported in behavioral and physiological studies in humans and other species [22-25]. The consistency between the results here and those in other species adds to growing evidence that the rat is a suitable animal model for mammalian binaural hearing in general.

The stimulus parameters investigated here were chosen for their relevance to binaural hearing in CI users. Factors which limit the ability of current human binaural CI users to discriminate ITDs are likely to include the poor synchronization of CI pulses to stimulus fine structure, and the fact that the pulse rate of CI processors are often quite high, running at 900 Hz or greater at each electrode channel. Despite the poor synchronization between the two ears under everyday listening conditions, some post-lingually deaf bilateral CI users do show good envelope ITD sensitivity (∼100 μs) at pulse rates at 100 Hz and sometimes up to 300 Hz [23, 26, 27]. However, sensitivity to rate discrimination or temporal information in both bilateral and unilateral CIs generally deteriorates dramatically at pulse repetition rates faster than about 300 pps [24, 28-31]. Our finding that neural sensitivity to ITD also declines sharply for acoustic pulse trains as pulse rates are increased beyond 300 Hz is thus consistent with these observations, and suggest that future animal studies testing different stimulus parameter types could help identify which possible changes to current stimulation strategies are likely to be most promising.

Our finding that Hanning windowed stimuli produced lower, but still substantial, neural sensitivity to ITD compared to rectangular windowed stimuli suggests that neural activity in the IC encodes both onset and ongoing ITD information. However, the IC’s ability to encode ongoing ITD cues of pulse train stimuli appears to decline dramatically when pulse rates reach or exceed 900 Hz. To improve ITD perception through CIs, the use of lower pulse rates would likely be preferable. Our data indicate that at rates of ∼300 Hz, neural and behavioral ITD sensitivity are high regardless of whether strong onset ITD cues are present. This is also consistent with human data showing that, for pulse rates above 200 Hz, both normally hearing [32] and CI listeners [30] tend to rely more heavily on onset ITD as opposed to ITD in the ongoing portion of sound. Even for CI users with exceptionally high sensitivity to ITD even at 600 Hz, sensitivity generally declined with the introduction of a 10 ms onset ramp [24].

Of course, spatial hearing is far from the only consideration that needs to be optimized in clinical CI processors, and there are theoretical reasons why some might expect high pulse rates to work better for encoding important aspects of speech sounds. However, experimental evidence usually fails to demonstrate any significant improvement in speech perception with pulse rates above 600 Hz [33]. Similarly, the expectation that faster pulse rates should improve the representation of temporal modulations also does not appear to supported by psychophysical evidence [34]. There seems to be no compelling evidence that pulse rates of ∼1000 Hz or more are necessary for important auditory tasks such as speech comprehension. Our data thus contribute to a growing body of evidence suggesting that using lower pulse rates in CIs could be beneficial, and our general approach shows that neural *d*′ measures may be a highly useful tool for predicting the likely effects of changing carrier pulse rates or stimulation patterns in CIs through animal studies.

### Ethics

All procedures were assessed and approved by the Animal Research Ethics Subcommittee of the City University of Hong Kong, and under license by the Department of Health of Hong Kong (Ref No.: (18-9) in DH/SHS/8/2/5 Pt.3).

## Data accessibility

The datasets and code supporting this article have been uploaded in Dryad Digital Repository: https://doi.org/10.5061/dryad.b5mkkwhcn [35].

## Authors’ contributions

K.L. carried out the behavioral training, electrophysiological recording, participated in the design of the study and drafted the manuscript; V.G.R carried out the data analysis and the statistical analyses, and critically revised the manuscript; A.P.M carried out the electrophysiological recording; C.H.K.C carried out the behavioral training; J.W.H.S conceived of the study, designed the study, coordinated the study, supervised the data analysis and the statistical analyses, and critically revised the manuscript. All authors gave final approval for publication and agree to be held accountable for the work performed therein.

## References

1. Seidl AH and Grothe B. Development of sound localization mechanisms in the mongolian gerbil is shaped by early acoustic experience. J Neurophysiol. 2005;94:1028–1036. (doi: 10.1152/jn.01143.2004)

2. Hancock KE, Noel V, Ryugo DK and Delgutte B. Neural coding of interaural time differences with bilateral cochlear implants: effects of congenital deafness. J Neurosci. 2010;30:14068–14079. (doi: 10.1523/JNEUROSCI.3213-10.2010)

3. Kan A and Litovsky RY. Binaural hearing with electrical stimulation. Hear Res. 2015;322:127–137. (doi: 10.1016/j.heares.2014.08.005)

4. Rosskothen-Kuhl N, Buck AN, Li K and Schnupp JW. Microsecond interaural time difference discrimination restored by cochlear implants after neonatal deafness. eLife. 2021;10. (doi: 10.7554/eLife.59300)

5. Litovsky RY, Jones GL, Agrawal S and van Hoesel R. Effect of age at onset of deafness on binaural sensitivity in electric hearing in humans. J Acoust Soc Am. 2010;127:400–414. (doi: 10.1121/1.3257546)

6. Li K, Chan CH, Rajendran VG, Meng Q, Rosskothen-Kuhl N and Schnupp JW. Microsecond sensitivity to envelope interaural time differences in rats. J Acoust Soc Am. 2019;145:EL341–EL347. (doi: 10.1121/1.5099164)

7. Luo L, Wang Q and Li L. Neural representations of concurrent sounds with overlapping spectra in rat inferior colliculus: Comparisons between temporal-fine structure and envelope. Hear Res. 2017;353:87–96. (doi: 10.1016/j.heares.2017.06.005)

8. Luo L, Xu N, Wang Q and Li L. Disparity in interaural time difference improves the accuracy of neural representations of individual concurrent narrowband sounds in rat inferior colliculus and auditory cortex. J Neurophysiol. 2020;123(2):695–706. (doi: 10.1152/jn.00284.2019)

9. Schnupp JWH, Garcia-Lazaro JA and Lesica NA. Periodotopy in the gerbil inferior colliculus: local clustering rather than a gradient map. Front neural circ. 2015;9:37. (doi: 10.3389/fncir.2015.00037)

10. Nelken I, Chechik G, Mrsic-Flogel TD, King AJ and Schnupp JW. Encoding stimulus information by spike numbers and mean response time in primary auditory cortex. J Comput Neurosci. 2005;19:199–221.

11. Shackleton TM, Skottun BC, Arnott RH and Palmer AR. Interaural time difference discrimination thresholds for single neurons in the inferior colliculus of guinea pigs. J Neurosci. 2003;23:716–724. (doi: 10.1523/JNEUROSCI.23-02-00716.2003)

12. Stecker GC, Harrington IA and Middlebrooks JC. Location coding by opponent neural populations in the auditory cortex. PLoS Biol. 2005;3:e78. (doi: 10.1371/journal.pbio.0030078)

13. Magezi DA and Krumbholz K. Evidence for opponent-channel coding of interaural time differences in human auditory cortex. J Neurophysiol. 2010;104:1997–2007. (doi: 10.1152/jn.00424.2009)

14. Młynarski W. The opponent channel population code of sound location is an efficient representation of natural binaural sounds. PLoS Comput Biol. 2015;11:e1004294. (doi: 10.1371/journal.pcbi.1004294)

15. Thompson SK, Von Kriegstein K, Deane-Pratt A, Marquardt T, Deichmann R, Griffiths TD and McAlpine D. Representation of interaural time delay in the human auditory midbrain. Nat Neurosci. 2006;9:1096–1098. (doi: 10.1038/nn1755)

16. Schnupp JWH, Hall TM, Kokelaar RF and Ahmed B. Plasticity of temporal pattern codes for vocalization stimuli in primary auditory cortex. J Neurosci. 2006;26(18):4785–95. (doi: 10.1523/JNEUROSCI.4330-05.2006)

17. Linsker R. Local synaptic learning rules suffice to maximize mutual information in a linear network. Neural Comput. 1992;4:691–702. (doi: 10.1162/neco.1992.4.5.691)

18. Chechik G. Spike-timing-dependent plasticity and relevant mutual information maximization. Neural Comput. 2003;15:1481–1510. (doi: 10.1162/089976603321891774)

19. Slee SJ and David SV. Rapid task-related plasticity of spectrotemporal receptive fields in the auditory midbrain. J Neurosci. 2015;35:13090–13102. (doi: 10.1523/JNEUROSCI.1671-15.2015)

20. Blackwell JM, Lesicko AM, Rao W, De Biasi M and Geffen MN. Auditory cortex shapes sound responses in the inferior colliculus. eLife. 2020;9. (doi: 10.7554/eLife.51890)

21. Walker KMM, Ahmed B and Schnupp JWH. Linking cortical spike pattern codes to auditory perception. J Cogn Neurosci. 2008;20(1):135–52. (doi: 10.1162/jocn.2008.20012)

22. Bernstein LR. Auditory processing of interaural timing information: new insights. J Neurosci Res. 2001;66:1035–1046. (doi: 10.1002/jnr.10103)

23. Laback B, Majdak P and Baumgartner W-D. Lateralization discrimination of interaural time delays in four-pulse sequences in electric and acoustic hearing. J Acoust Soc Am. 2007;121:2182–2191. (doi: 10.1121/1.2642280)

24. van Hoesel RJ, Jones GL and Litovsky RY. Interaural time-delay sensitivity in bilateral cochlear implant users: Effects of pulse rate, modulation rate, and place of stimulation. J Assoc Res Otolaryngol. 2009;10:557. (doi: 10.1007/s10162-009-0175-x)

25. Chung Y, Hancock KE and Delgutte B. Neural coding of interaural time differences with bilateral cochlear implants in unanesthetized rabbits. J Neurosci. 2016;36:5520–5531. (doi: 10.1523/JNEUROSCI.3795-15.2016)

26. van Hoesel RJ and Tyler RS. Speech perception, localization, and lateralization with bilateral cochlear implants. J Acoust Soc Am. 2003;113:1617–1630. (doi: 10.1121/1.1539520)

27. Van Hoesel RJ. Sensitivity to binaural timing in bilateral cochlear implant users. J Acoust Soc Am. 2007;121:2192–2206. (doi: 10.1121/1.2537300)

28. Shannon RV. Multichannel electrical stimulation of the auditory nerve in man. I. Basic psychophysics. Hear Res. 1983;11:157–189. (doi: 10.1016/0378-5955(83)90077-1)

29. Townshend B, Cotter N, Van Compernolle D and White R. Pitch perception by cochlear implant subjects. J Acoust Soc Am. 1987;82:106–115. (doi: 10.1121/1.395554)

30. van Hoesel RJ. Observer weighting of level and timing cues in bilateral cochlear implant users. J Acoust Soc Am. 2008;124:3861–3872. (doi: 10.1121/1.2998974)

31. Venter PJ and Hanekom JJ. Is there a fundamental 300 Hz limit to pulse rate discrimination in cochlear implants?. J Assoc Res Otolaryngol. 2014;15:849–866. (doi: 10.1007/s10162-014-0468-6)

32. Stecker GC and Hafter ER. Temporal weighting in sound localization. J Acoust Soc Am. 2002;112:1046–1057. (doi: 10.1121/1.1497366)

33. Shannon RV, Cruz RJ and Galvin JJ. Effect of stimulation rate on cochlear implant users’ phoneme, word and sentence recognition in quiet and in noise. Audiol Neurootol. 2011;16:113–123. (doi: 10.1159/000315115)

34. Green T, Faulkner A and Rosen S. Variations in carrier pulse rate and the perception of amplitude modulation in cochlear implant users. Ear Hear. 2012;33:221–230. (doi: 10.1097/AUD.0b013e318230fff8)

35. Li K, Rajendran VG, Mishra AP, Chan CH and Schnupp JWH. Data from: Data_IC_ITD_dprime_study. Dryad Digital Repository. 2021. (doi: 10.5061/dryad.b5mkkwhcn)

